# Critical flicker fusion thresholds predict attentional blink magnitude

**DOI:** 10.64898/2026.06.10.731301

**Authors:** Clinton S. Haarlem, Jake G. Tiernan, Maggie Kelly, Lochlan Cooney, Andrew L. Jackson, Kevin J. Mitchell, David P. McGovern, Redmond G. O’Connell

## Abstract

The critical flicker fusion (CFF) threshold is a psychophysical measure used to quantify the temporal resolution of the visual system and is known to vary across individuals. However, it is unclear if this measure is stimulus-specific, or if it may represent a more fundamental processing rate for visual perception in general. Here, we assess if individual variation in CFF is predictive of two features of visual processing that are dependent on temporal perception: the attentional blink and global motion sensitivity. In a non-clinical sample of 84 individuals, flicker fusion thresholds were predictive of the magnitude of the attentional blink. In contrast, we found no link between flicker fusion and global motion sensitivity in a sample of 79 individuals. Our results suggest that CFF reflects a visual processing rate that impacts other, more complex perceptual tasks.

## Introduction

The critical flicker fusion threshold (CFF) is a psychophysical measure used to quantify the temporal resolution of the visual system (Brozek & Keys, 1945; Landis, 1954). It is defined as the point at which a luminous stimulus flickers at such a high rate that the flicker can no longer be perceived and the stimulus appears steady (Hecht & Verrijp, 1933; Misiak, 1947). CFF has been a topic of scientific study for many decades, yet its role in overall visual perception is still poorly understood. Recent cross-species investigations have established that CFF systematically tracks ecological niche such that species that inhabit fast paced sensory environments exhibit higher CFF (Boström et al., 2016; Haarlem et al., 2026; Potier et al., 2020). In humans, CFF appears to be a stable trait that varies substantially across individuals in the general population (Haarlem et al., 2024) and is sensitive to certain pathologies and altered nervous system states (Curran & Wattis, 2000; Hindmarch, 1982; Kahlbrock et al., 2012; Mark et al., 1958; Smith & Misiak, 1976). While the inter-species and inter-individual variations that have been observed are of a magnitude that seem likely to impact perceptual performance (e.g. 30 Hz range across individuals in Haarlem et al., 2024), it is unclear which perceptual tasks depend on CFF. Visual perception is hierarchically organised (Felleman & Van Essen, 1991), and CFF requires only the detection of changes in luminance; a low-level aspect of visual processing. As such, it is unclear whether temporal resolution estimates based on CFF are stimulus-specific or reflect a more fundamental visual processing rate that impacts how more complex stimuli are processed. To date, the impact on perceptual performance of the considerable variation in CFF found in humans has not been systematically tested.

If the CFF threshold is representative of how much visual information an individual can process per unit time, then it should predict performance on tasks incorporating other types of rapidly appearing visual information. One commonly studied example of such a task is the rapid serial visual presentation (RSVP) task (Lawrence, 1971) in which participants are required to identify target stimuli appearing within a series of rapidly presented distractor stimuli. One of the most extensively studied applications of RSVP is the attentional blink (AB), a phenomenon which occurs when a participant is tasked with identifying two targets appearing in close temporal proximity to each other (Raymond et al., 1992). Although the first target is reliably detected, the second target is frequently missed if it appears within approximately 500 ms of the first (Arnell et al., 2006; Broadbent & Broadbent, 1987; Chun & Potter, 1995; Dux & Marois, 2009; Raymond et al., 1992). While numerous theories have been proposed regarding the specific mechanisms underlying the AB, behavioural and neurobiological studies indicate that the phenomenon is related to the rate with which attention can be allocated to a visual target (Dux & Marois, 2009; Martens & Wyble, 2010; Raymond et al., 1992; Shapiro & Raymond, 2025). We hypothesize that if CFF is indicative of a baseline processing rate for overall visual perception, then individuals with higher CFF should encode visual information faster, thus reducing the AB effect.

Another perceptual function in which CFF could plausibly play a role is in the perception of motion. CFF is higher in species for which high-speed motion perception is crucial to survival (Boström et al., 2016; Haarlem et al., 2026; Potier et al., 2020). For example, faster, more manoeuvrable species of bird possess higher CFF than closely related, slower species (Boström et al., 2016; Potier et al., 2020). Neuroimaging studies have also shown that there is significant overlap in the areas of the visual cortex that process flicker and motion stimuli, suggesting the two types of stimuli may be tightly linked (Kolster et al., 2010; Mullen et al., 2010; Spitschan et al., 2016). Recent work using the random-dot-motion (RDM) direction discrimination task has also shown that raising the stimulus display frame rate leads to increased misperception of the direction of motion (Mc Keown et al., 2023). We therefore also hypothesize that higher CFF thresholds are linked with a higher sensitivity to motion energy, resulting in an increased ability to discriminate motion direction.

Here we test these hypotheses by examining whether individual variation in flicker fusion thresholds predicts the magnitude of the AB effect, as measured through the RSVP task, and motion discrimination sensitivity, as measured through the RDM task. Our rationale is that if CFF reflects a stimulus-specific threshold, it should bear little relation to performance on either task. If, however, it indexes a more fundamental rate of visual processing, higher CFF thresholds should predict a smaller attentional blink magnitude, as well as a higher sensitivity to motion, allowing for increased direction discrimination ability.

## Methods

All experimental procedures were conducted in accordance with the Declaration of Helsinki and EU GDPR. Experimental sessions were conducted during office hours (9am – 5pm) in a darkened, windowless booth. Before the start of the experiment a short dark adaptation phase took place during which the experimenter explained the three tasks to the participant. Data collection was carried out by LC, MK and CH.

### Participants

We recruited 91 participants, 64 females (mean age 20.61, SD = 1.98) and 25 males (mean age 20.88, SD = 2.68). Two participants identified as neither male nor female (ages 20 and 24). Participants consisted of students and research staff recruited from Trinity College Dublin, as well as personal acquaintances of the experimenters. All participants provided written, informed consent and all participants had normal or corrected-to-normal visual acuity. Participants were compensated either with a 10 Euro e-voucher, with teaching credits forming part of their postgraduate research degree (at the rate of 1 credit per half-hour of participation) or participated on a voluntary basis.

One participant was excluded due to failure to obtain a consistent CFF measurement. Six participants were excluded from RSVP analysis: two participants were excluded due to the researcher’s usage of erroneous stimulus timings and for four participants the data failed to save. 11 participants were excluded from the RDM analysis: eight participants were unable to reach the desired accuracy range of 60 - 80% (see RDM methods below) during the session, and for three participants the data failed to save. Thus, 84 participants were included in the RSVP analysis and 79 participants were included in the RDM analysis.

During model diagnostics, two data points corresponding to the participants who identified as neither male nor female were flagged with high hat values and large Cook’s distance values, indicating that these points were highly influential and had high leverage. However, analysis of studentized residuals indicated that these two participants were not statistical outliers. For these reasons, we analysed the data both with and without these participants included. Below, we report all statistical outputs with these participants included, but flag any instances where their exclusion altered the statistical conclusions.

### CFF

CFF thresholds were measured using a variation of a device used in previous research (Haarlem et al., 2024a; Haarlem et al., 2024b). The device consisted of a black box housing a standard, white, 5mm LED, operated with an Arduino Uno R3 microprocessor. Mean luminance of the LED was ~ 282 cd/m^2^. The housing included a 16 cm long viewing tube through which participants observed the LED, standardising the viewing distance (see supplementary material S1). To minimise ambient light, participants were asked to wear lensless goggles with a blacked-out frame during measurement. During testing, the device was kept on top of the desktop computer and participants were instructed to adjust their seat height before the start of the experiment so that they could comfortably look through the viewing tube.

We employed the psychophysical method of adjustment to measure CFF (Fig 1, top). Participants used a rotary encoder dial to slowly decrease flicker frequency of the LED in 1 Hz increments until flicker first became visible, which was indicated verbally to the experimenter. We took five measurements per participant, with starting frequencies in the following order: 90, 95, 90, 90 and 95 Hz. Participants were not informed of the differing starting values. We took the mean of the five measurements as the participant’s CFF threshold. Although the method of adjustment typically involves averaging descending and ascending measurements, we opted to employ only descending trials, as previous research showed lower test-retest reliability for ascending trials (Haarlem et al. 2024a). Participants were given a short practice run prior to the measurements to allow them to familiarise themselves with the device.

**Figure 1.**
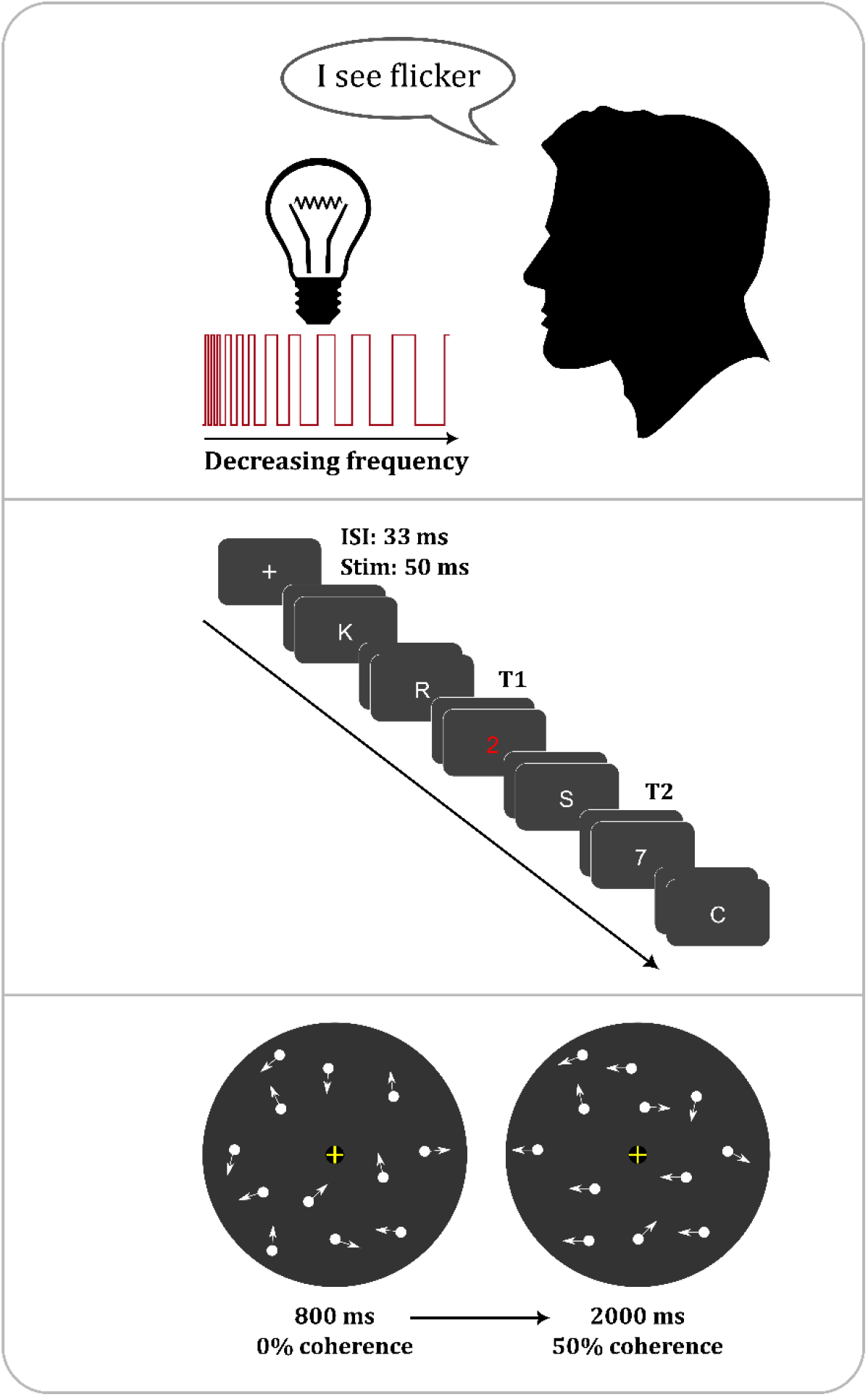
Overview of psychophysical tasks. Participants started with the CFF task, measured with the descending method of adjustment (top), followed by the RSVP task (middle). The RSVP panel shows a schematic overview of a lag 2 trial. White letters were used as distractor stimuli, a red single digit was used as T1 and a white single digit was used as T2 (note that the actual number of stimuli used in each trial was 18). Empty frames behind distractors represent ISI. The RDM task (bottom) was completed last for each participant. RDM panel shows a schematic overview of a trial on the RDM task (excluding lead-in period). Arrows indicate movement of each white dot within a large round aperture.

### RSVP - attentional blink

The RSVP task was created in MATLAB (The Mathworks, 2022) and Psychtoolbox-3 (Kleiner et al., 2007) and was modelled after similar tasks used in previous research (Arnell & Larson, 2002; Arnell et al. 2006; Jefferies et al., 2008; Thomson et al., 2015). The task was displayed on a 1920×1080 resolution monitor with a 60 Hz refresh rate. During a trial, participants were shown a stream of 18 stimuli, consisting of 16 capitalised distractor letters, a target digit (T1) and a probe digit (T2), presented in Arial font (Fig 1, middle). The stream of distractors consisted of a random combination of non-repeating letters, excluding “B”, “I”, “O” and “S”, to avoid potential confusion with similar looking digits. T1 and T2 were randomly selected per trial from single digits 0-9, with the conditions that T1 and T2 could not be the same. Stimuli subtended ~ 1 degree of visual angle and participants were seated at a viewing distance of ~ 60 cm. The distractors and T2 were always presented in white, T1 was always presented in red.

Each stimulus appeared for 50 ms, with a 33 ms interstimulus interval, for a total stream length of 1.5 secs. T1 appeared randomly in the stream, but was always preceded by a minimum of six distractors. Following T1, T2 appeared either directly after T1 (lag 1), after a single distractor (lag 2), after two distractors (lag 3) or after several additional distractors (lag 4 – lag 7). Following the stimulus stream, participants were required to type which two digits were shown, in the correct order, using a keyboard. After a practice block of 10 trials, participants completed two blocks of 70 trials, resulting in a total of 20 trials for each lag condition.

### RDM

In the RDM task, participants viewed a cloud of dots that initially moved randomly. After a fixed lead-in period, a proportion of the dots started to move coherently in a common direction while the remainder continued to move in random directions, and participants were required to indicate the direction of coherent motion. The RDM task is commonly used to assess an individual’s sensitivity to global motion, by varying the percentage of coherently moving dots (Newsome & Pare, 1988; Britten et al., 1992). The version of this task used in the current study was created during previous research conducted by our research group (Grogan et al., 2023; Grogan et al., 2025) and was run using Matlab (The Mathworks, 2022) and Psychtoolbox-3 (Kleiner et al., 2007). The task was displayed using the same set-up as for the RSVP task. During a trial, a central fixation cross and a white ring (10° diameter) appeared on a grey background for 400 ms, after which a cloud of 60 randomly moving dots, 0.15° diameter, appeared within an 8° diameter aperture. After 800 ms, a percentage of the dots started moving, either to the left or right with equal probability across trials (Fig 1, bottom). Coherently moving dots were displaced relative to their position three frames earlier, at a speed of ~18° per second, with the remaining dots displaced randomly. Participants responded to the trial with a keyboard press, using either ‘C’ if they perceived leftward motion or ‘N’ if they perceived rightward motion. Trials had a deadline of 2000 ms and if participants failed to respond before the deadline, the response was recorded as incorrect. The fixation cross momentarily changed either to green after a correct response or to red after an incorrect response to provide participants with feedback after each trial. The motion direction discrimination task consisted of four parts: first, participants completed a practice block with dot coherence level set to 80%. The block consisted of 30 trials but ended early if participants responded correctly on 10 consecutive trials. Second, participants performed a block which gradually increased in difficulty: coherence level started at 50% and then decreased to 40%, 30%, 25%, 20%, 15% and finally 10%.

Participants completed eight consecutive trials before each increase in difficulty. After this block participant performance was evaluated and the coherence level that most closely matched an accuracy with an upper bound of 70% was set as the starting coherence level for the next step. The third step consisted of one block of 60 trials using the Quest procedure (Watson & Pelli, 1983). This procedure actively titrated coherence level to achieve an accuracy of ~70%. Lastly, participants performed a final block of 30 trials using the suggested coherence level resulting from the Quest procedure to verify participants were performing within 60 – 80% accuracy. If performance fell outside of this range, the Quest procedure was repeated with a manually adjusted starting value.

### Statistical analysis

We extracted raw performance scores and pre-processed the task data using custom MATLAB scripts. We carried out the main statistical analyses using R version 4.5.2 (R Core Team, 2025) and RStudio version 2026.01.0.392 (Posit team, 2026). We used functions from the “car” package (Fox & Weisberg, 2019) for model diagnostics.

#### RSVP & Attentional blink

We assessed the relationship between CFF and T1 accuracy by creating a linear regression model using CFF as predictor and T1 accuracy as response variable.

For an attentional blink to occur, it is necessary for T1 to be successfully attended prior to the occurrence of T2 (Chun & Potter, 1995; Potter et al., 1998). Therefore, to assess the attentional blink effect we only analysed trials for which T1 was correctly identified. To examine the effect of CFF on the attentional blink, we calculated three different metrics for each participant. First, we calculated AB-magnitude by subtracting mean T2 accuracy on the lag for which group-level accuracy was lowest (lag 4) from mean accuracy on the latest lag (lag 7). We then created binomial regression models for overall individual performance on T2, using the lag condition as the predictor and accuracy as the response variable. For each participant, the intercept for the respective model was taken as a measure for baseline performance on T2 across all lag levels, and the slope was used as a measure for the rate of recovery from the AB effect across lag levels. Using all three AB metrics as response variables, we conducted a multivariate regression to assess the effect of CFF on the overall profile of the attentional blink effect. As each of the three response variables had a different magnitude, we standardized them in the multivariate models by using the ‘scale’ function in R, which for each response variable subtracts the mean and divides by the standard deviation. We included age as a continuous variable and gender as a three-level factor (female, male, other) in both the T1-accuracy and AB models as additional predictor variables.

For multivariate regression, it is common practice to follow up on statistically significant models with a post-hoc inspection to identify which response variable(s) may have the largest effect on the model. In this study we employ the post-hoc Roy-Bargmann stepdown procedure (Roy, 1958; Roy & Bargmann, 1958; Tabachnick & Fidell, 2013), consisting of a set of univariate linear regression models. For this procedure, the priority of the response variables is determined a priori. The first priority response variable is modelled first. If this model shows a significant effect, the second priority response variable is modelled, with the first response variable included as a covariate. This process continues until a non-significant model is produced. We have determined the priority of our response variables as follows:

1. AB magnitude. Our main hypothesis is that individuals with higher CFF thresholds will exhibit a reduced attentional blink effect.
2. Overall T2 accuracy (Intercept). We theorize that overall accuracy across all T2 lags is dependent on many factors such as task familiarity, fatigue level, sustained attention etc., and thus would be predicted by CFF to a lesser degree than AB magnitude.
3. AB recovery rate (Slope). We theorize that since recovery rate is directly dependent on both other response variables (the maximum amount of potential recovery is limited by both the magnitude of the AB and overall performance), this response variable will capture the least amount of unique variance in the data.

#### RDM

We examined the possible effect of CFF on global motion sensitivity by extracting the lowest percentage of coherently moving dots at which a participant still performed within an accuracy range of 60 – 80 %. We used linear multiple regression, with RDM coherence threshold as the response variable and CFF, age and gender as predictors. During data collection, one of the experimenters employed a slightly different protocol from the other two if an initial verification block of trials at the suggested coherence level indicated by Quest did not result in an accuracy range of 60 – 80%. When this situation occurred, two of our experimenters had participants redo the Quest block for a second titration, followed by another verification run. Our third experimenter manually adjusted dot coherence and reran the verification block until accuracy was within the desired range. We analysed the data both with the two differing methodologies combined into a single dataset, as well as with them separated. Overall results did not differ, thus in the Results section we only report the results for the combined dataset. Results of the separate analysis are available in the supplementary material (S2).

## Results

### RSVP task

#### T1 performance

We first examined RSVP task performance beginning with T1 accuracy to confirm that participants successfully attended the first target before turning to T2 performance and the attentional blink. Mean T1 accuracy was high across all T2 lags (mean = 0.91, standard deviation (SD) = 0.06) with the exception of lag 1, where mean accuracy fell to 0.61 (SD = 0.17). This reduced T1 performance for lag 1 was accompanied by relatively high accuracy for T2. The latter is commonly observed on RSVP tasks and is referred to as lag 1 sparing (Potter et al., 1998, see Discussion).

We conducted a linear regression with T1 accuracy as the response variable and found no effect of CFF (β = 0.00, P = 0.686,) or age (β = 0.00, P = 0.939, model r^2^ = 0.01). Males had significantly higher T1 accuracy (β 0.03, P = 0.038), with a mean accuracy of 0.93 (SD = 0.04), compared to females, who had a mean T1 accuracy of 0.90. Mean accuracy of participants who identified as neither male nor female was 0.93 (SD = 0.05) did not differ significantly from the female reference (β = 0.27, P = 0.511).

#### Attentional blink

Examining T2 accuracy as a function of lag confirmed that our participants exhibited the classic attentional blink effect with accuracy reducing for lags 2, 3 and 4 and then recovering for later lags (Fig 2). In addition, participants exhibited the lag 1 sparing effect typically observed in the attentional blink literature (Chun & Potter, 1995; Dux & Marois, 2009, Potter et al., 1998; Potter et al., 2002).

**Figure 2.**
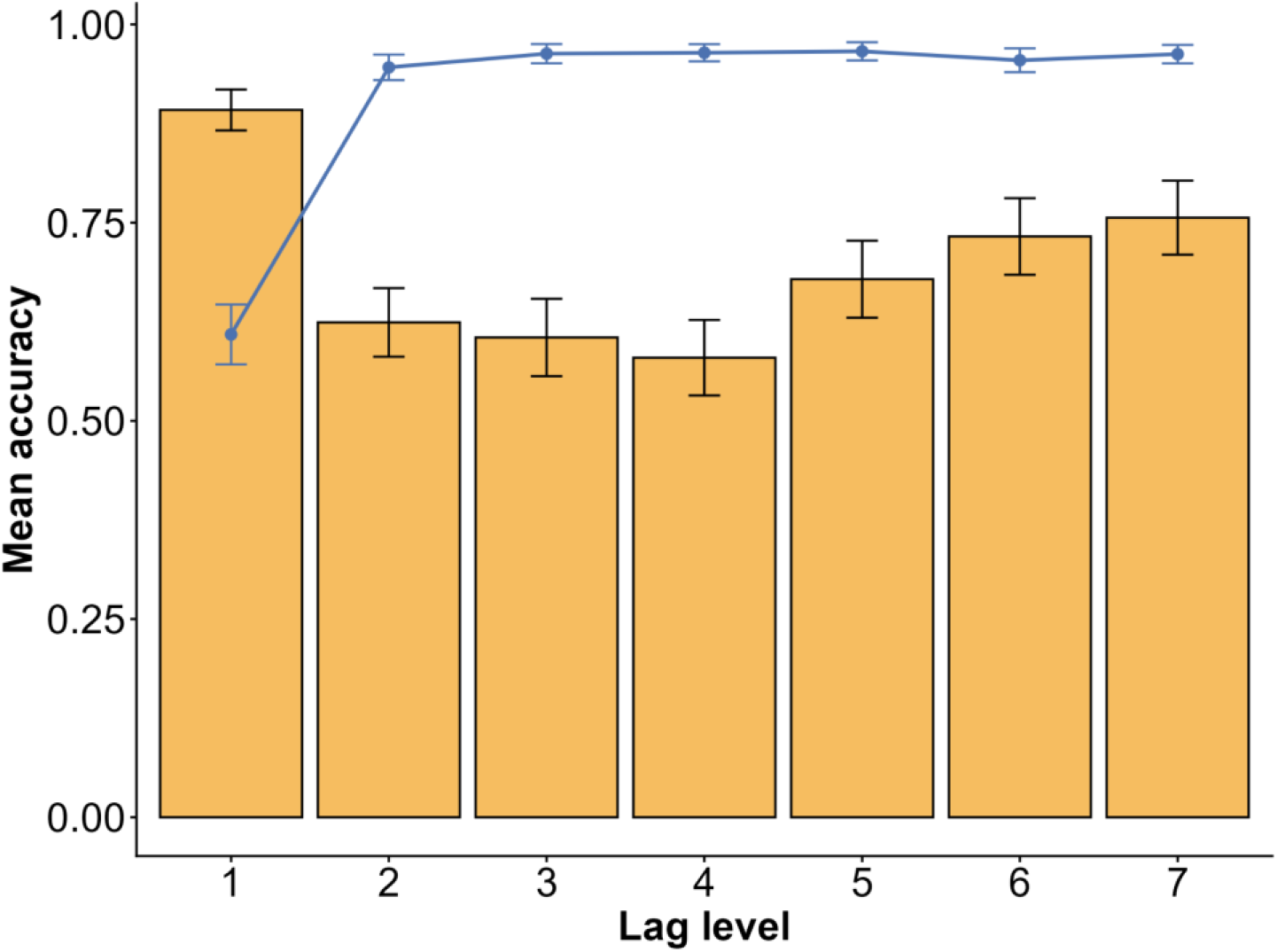
Attentional blink effect. Group level mean accuracy is shown across all T2 lag levels for T1 (connected dots) and T2 (bars). Error bars indicate standard error of the mean. Lag levels 1-7 correspond to SOA’s of 83, 167, 250, 333, 417, 500 and 583 ms respectively.

A multivariate regression with the overall AB-profile as the response variable revealed a significant effect of CFF (Pillai’s trace 0.11, P = 0.029) and gender (Pillai’s trace 0.16, P = 0.047), but not age (Pillai’s trace = 0.05, P = 0.239). Although gender showed the largest effect in this model, removal of the two participants which were flagged for high leverage and high influence as outlined in the Methods section, changed this result. With these participants excluded, CFF became the strongest predictor of AB magnitude (Pillai’s trace = 0.12, P = 0.018), followed by gender (Pillai’s trace = 0.12, P = 0.022), with age remaining non-significant (Pillai’s trace = 0.08, P = 0.113). Observation of the scatter plots and lines of best fit in Figure 3 indicated that participants with higher CFFs had reduced AB-magnitudes and higher overall T2 accuracy but a reduced AB recovery rate.

**Figure 3.**
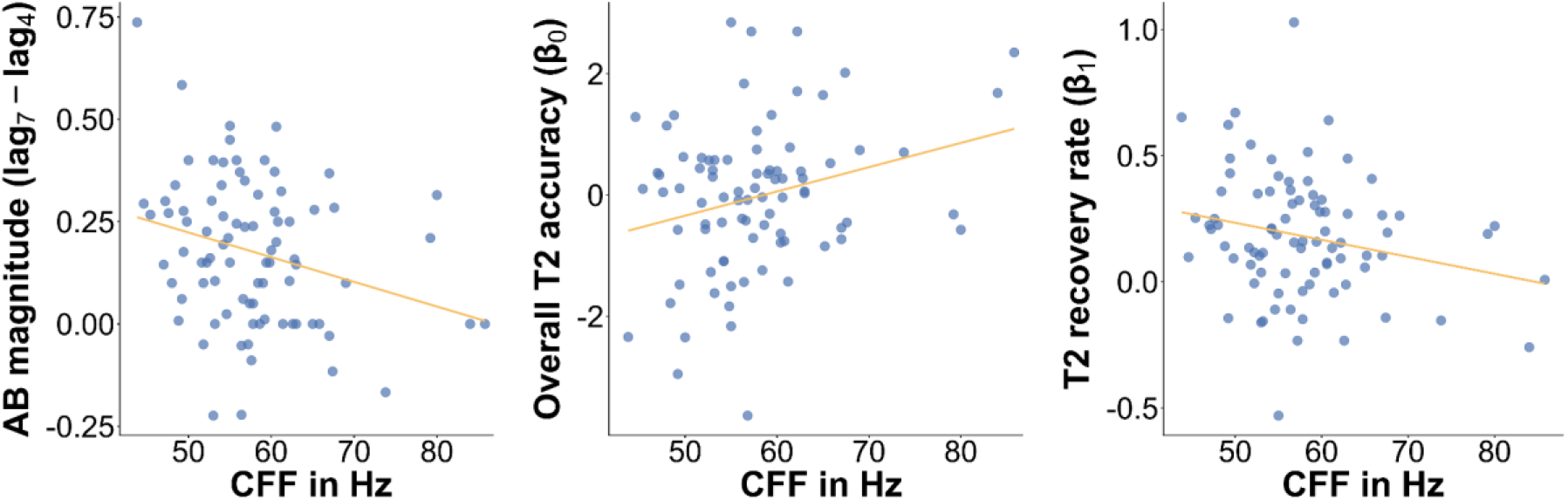
CFF and the AB-profile. Relationships between CFF and AB magnitude, as measured by subtracting lag 4 from lag 7 (left), overall T2 accuracy across all lags, as quantified by the intercept estimate (β_0_) of individual performance modelling (middle) and recovery rate from the AB effect across lags, as quantified by the slope estimate (β_1_) of individual performance modelling (right).

To follow up the significant multivariate effect, we conducted a Roy-Bargmann stepdown procedure with AB magnitude entered as the first priority response variable. AB magnitude accounted for most of the variance in the model, with CFF emerging as the only significant predictor; neither gender nor age were significant. We did not observe a significant effect of CFF or age in the follow-up model testing our second priority response variable, overall T2 accuracy. However, in this model we observed a significant effect of gender, with males having higher T2 accuracy than females (β = 0.83, P = 0.002). The participants who did not identify as male or female did not differ significantly from females (β = 0.24, P = 0.738). For full model outputs of the post-hoc analysis, see supplementary material S3.

To gain more insight into the relationship between CFF and AB, we separated our participants into high- and low CFF groups based on a median split (N = 42 for each group) and plotted T1 and T2 accuracy across all lag levels for each group (Figure 4). Figure 4 shows that the two groups show similar T1 accuracy and converge to similar T2 accuracy by lag 7, but differ markedly in AB magnitude (see lags 2-4) This figure also helps to account for the unexpected negative relationship between CFF and AB recovery rate indicated in Figure 3: the shallower recovery slope of the high CFF group reflects their smaller AB-magnitudes, rather than a slower return to baseline.

**Figure 4.**
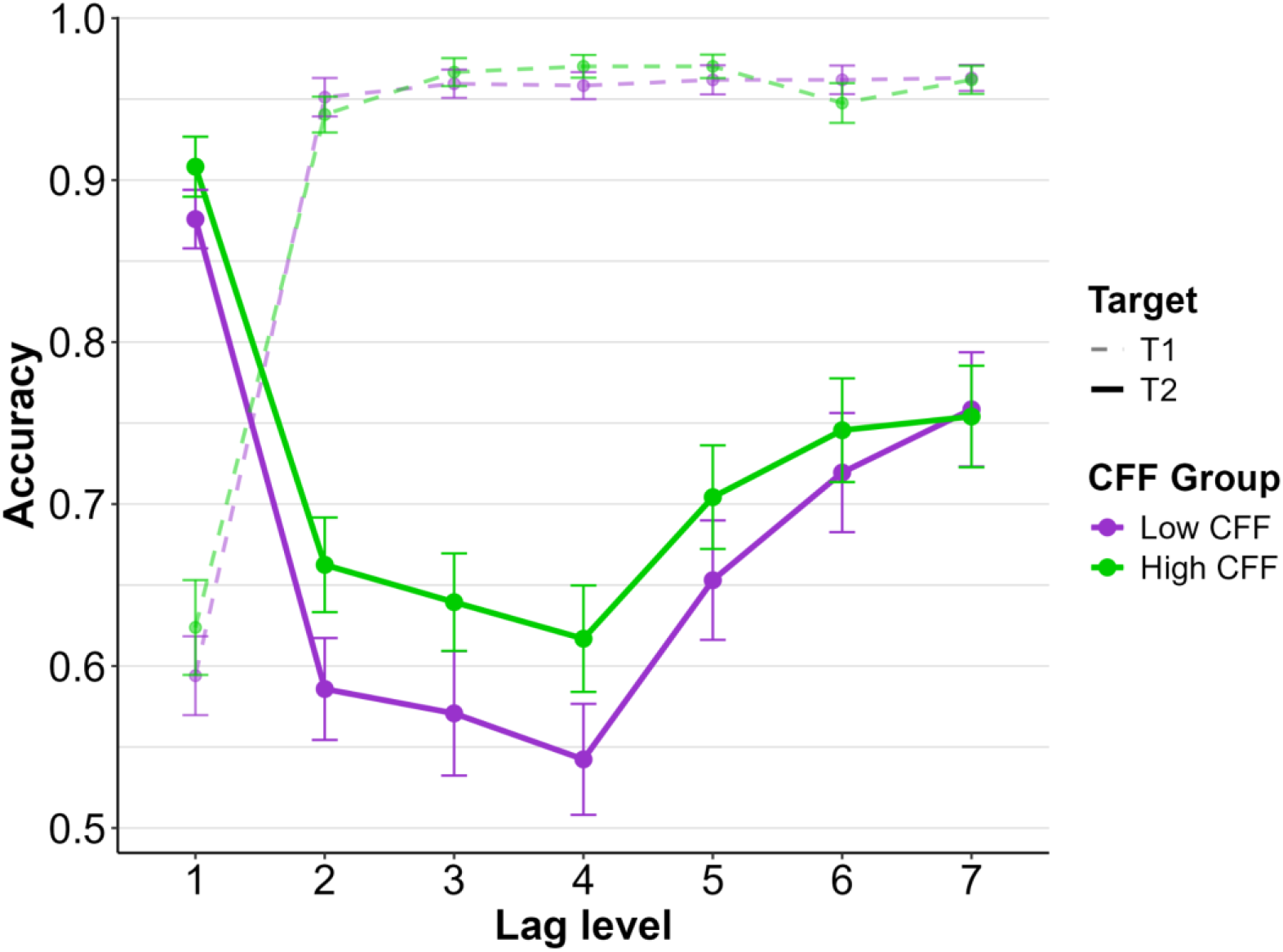
Group comparison of RSVP accuracy across all lag conditions. The high CFF group (green solid line) shows a smaller AB magnitude and higher overall T2 accuracy across all lags, compared to the low CFF group (purple solid line). No difference is observed for T1 accuracy (dashed lines). Lag levels 1-7 correspond to SOA’s of 83, 167, 250, 333, 417, 500 and 583 ms respectively. Points indicate group level mean accuracy, error bars indicate standard error of the mean.

### RDM task

We next examined whether CFF was associated with performance on the RDM task. As with the RSVP analyses, we modelled motion coherence thresholds as a function of CFF, age and gender. While participants with higher CFFs tended to show lower coherent motion thresholds, this relationship was not statistically significant. The overall model did not explain a significant proportion of the variance in motion coherence thresholds (adjusted r^2^ = −0.01, P = 0.557) and none of CFF, age and gender emerged as significant predictors (Table 1, Fig. 5).

**Table 1.** Linear model results for RDM coherence threshold. Estimates and p-values for each of our predictor’s effects on the response variable RDM coherence threshold.

| Predictor | Estimate ($\beta$) | P-value |
| --- | --- | --- |
| CFF | -0.19 | 0.419 |
| Age | -1.09 | 0.226 |
| Gender (male) | 5.01 | 0.253 |
| Gender (other) | 0.94 | 0.955 |

**Figure 5.**
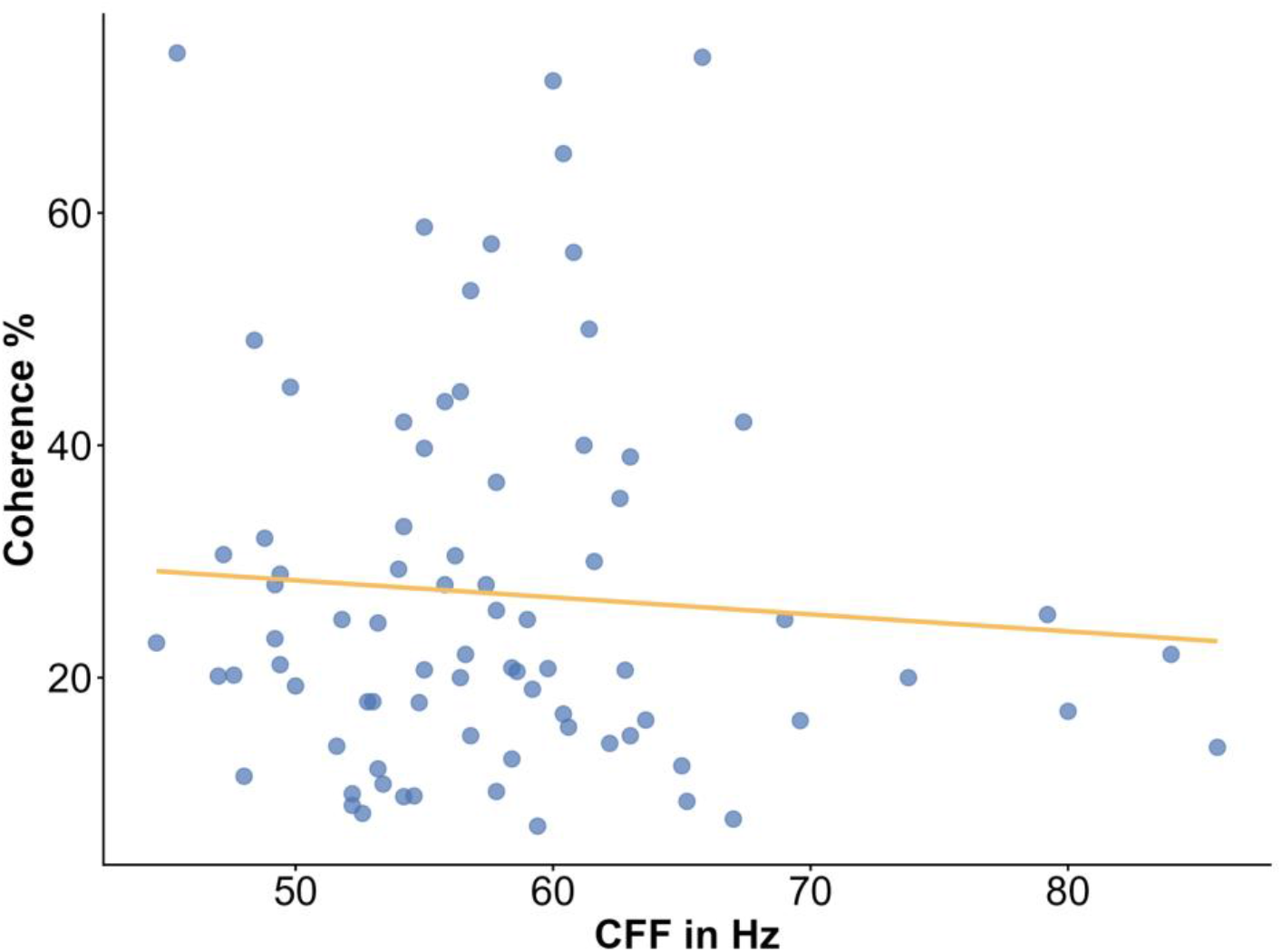
RDM coherence and CFF. The lowest coherence percentage at which a participant was able to perform with 60 - 80% accuracy on the RDM task (y-axis), plotted against CFF threshold (x-axis).

## Discussion

To test the theory that CFF measures a fundamental perceptual processing rate for the visual system, we examined the degree to which individual differences in CFF predicted performance on two different psychophysical measures of visual perception ability. Our results reveal that CFF predicts the magnitude of the attentional blink (AB) during rapid-serial-visual-presentation (RSVP) task performance, but there was no relationship between CFF and motion coherence thresholds as measured with the random-dot-motion task. To our knowledge this is the first study to examine the effects of individual variation in CFF on perceptual performance.

### RSVP task

#### T1 identification

Aside from using the RSVP task to probe the AB, we were also interested to know whether individual differences in CFF would predict T1 accuracy. Here, we observed no such effect. Detection of a single target is a relatively easy feat for the human visual system (Dux & Marois, 2009; Potter et al., 2002), and previous literature shows that individuals can even identify complex full visual scenes such as photographs in RSVP streams with presentation rates as fast as 13-17ms (Broers et al., 2018; Mohsenzadeh et al., 2018; Potter et al., 2014). Thus, considering the high accuracy on T1 across all lags, the lack of an influence of CFF in the current study is likely due to a ceiling effect.

#### Attentional blink

In contrast to T1 detection, we did observe a link between CFF and the AB profile. Although our statistical model included overall T2 accuracy (calculated as the intercept in the regression models of individual T2 performance across all lags), AB recovery rate (calculated as the slope in the regression models of individual T2 performance across all lags) alongside AB-magnitude, our post-hoc analyses indicated that most of the variance in the data was explained by AB magnitude. Although the post-hoc stepdown procedure did not find CFF to have a statistically significant effect on overall T2 accuracy, Figure 4 shows that the high CFF group did have slightly higher accuracy on T2 on all but the longest lag condition compared to the low CFF group. Additionally, the high CFF group had higher accuracy than the low CFF group on both T1 and T2 for the lag 1 condition. These findings suggest that CFF is not stimulus-specific and that the measure may indeed be indicative of a general visual processing rate.

Our results suggest that CFF is not strongly linked with the rate with which individuals recover from the attentional blink. However, it is important to note that the AB is normally present for around 500 ms after presentation of T1 (Arnell et al., 2006; Broadbent & Broadbent, 1987; Dux & Marois, 2009; Raymond et al., 1992). In our study, all but the longest lag condition fall within this window, which may explain why accuracy on T2 did not return to T1 baseline level even at lag 7 (Fig. 2). Thus, it is possible we did not manage to capture the full extent of the AB effect with this study and we are therefore unable to assess how CFF may influence recovery rate throughout the full timeframe required to return to baseline performance level.

#### Lag 1 effects

Consistent with most previous AB studies, we observed the lag 1 sparing effect in which T2 accuracy is high if it immediately follows T1. However, in our study, T1 accuracy at lag 1 fell well below T2 accuracy, a pattern less commonly reported in letter/digit AB studies. Interestingly, a similar effect was also observed in Potter et al. (2002), specifically at short stimulus onset asynchronies. The authors theorized that when T2 directly follows T1 with a very short SOA, the two stimuli may compete for processing at a relatively early stage of processing. As the main AB effect is thought to be linked with attention allocation during later stages of processing, this phenomenon occurring only at lag 1 may be due to a separate mechanism.

### RDM task

We found no correlation between CFF and motion coherence thresholds in the RDM task. Several factors may account for this null result. First, the RDM task is known to have a high initial difficulty for naive participants, often requiring a substantial amount of practice before performance stabilises (Corbett et al., 2023; Saffel & Matthews, 2003). As we were limited to a single session, we deemed it necessary to employ a relatively wide range for acceptable accuracy in order to maintain a viable sample.

However, this may have introduced additional measurement noise, which could have obscured a correlation with CFF if the effect is relatively small. Second, the RDM task may not capture the aspect of motion processing most likely to be impacted by variation in CFF. The RDM task indexes sensitivity to *global* motion: as each coherently moving dot’s trajectory is based on its position relative to three frames prior, it is impossible to extract motion direction by tracking any individual dot. An observer can only obtain a sense of global motion when perceiving the full stimulus field. Global motion perception relies on integration mechanisms in area MT/V5 and on decision processes operating on noisy evidence (Britten et al., 1992; Newsome & Paré, 1988), neither of which maps directly onto the temporal-resolution component that CFF is thought to index. Thus, although our results indicate that CFF does not strongly predict performance on this type of motion discrimination, it is very possible that CFF does show associations with other, lower level motion tasks that more directly index the temporal limits of motion perception. For example, variation in CFF may predict performance on tasks that measure duration thresholds for motion direction or speed discrimination (Burr, 1981; Snowden & Braddick, 1991) or temporal frequency thresholds for drifting stimuli (Burr & Ross, 1982). Because these measures assess how briefly or rapidly a moving stimulus can be modulated before temporal resolution breaks down, they are arguably more similar conceptually to the CFF and therefore should show stronger associations with individual differences in CFF

In conclusion, our results show that individual variation in CFF thresholds predicts the magnitude of the attentional blink but not performance on a motion discrimination task. This dissociation suggests that the CFF does not only indicate a temporal processing limit for low-level visual information such as the presence or absence of light, but may generalize to the processing of higher order information and the identification of distinct visual targets.

## Research Transparency Statement

### General disclosures

#### Artificial intelligence

Microsoft Copilot was used for the debugging of task code. No other AI assisted technology was used for this study or for the creation of this article.

#### Ethics

The study was approved by the Ethics Committee of Trinity College’s School of Psychology (ref: 4907).

### Study disclosures

#### Preregistration

No aspects of this study were preregistered.

#### Materials

A custom CFF measuring device was used, which is described in the Methods section. The RSVP- and RDM tasks were created in MATLAB. Task and preprocessing scripts are available at 10.5281/zenodo.20544462.

#### Data

Raw data files are available at 10.5281/zenodo.20544085.

#### Analysis scripts

Statistical analysis scripts are available at 10.5281/zenodo.20544085.

