## Supplementary material for "Critical flicker fusion thresholds predict attentional blink magnitude"

**S1: Schematic of CFF measuring tool**

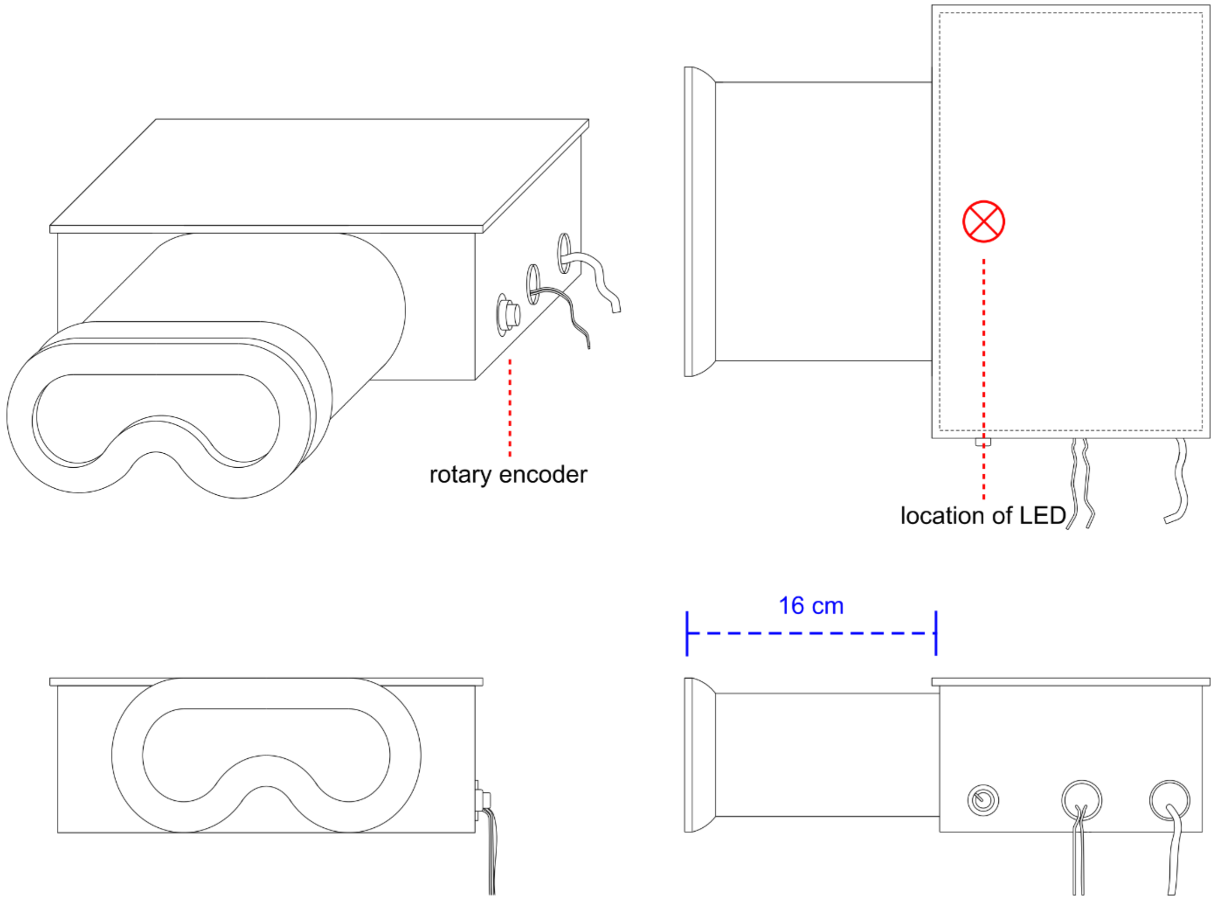

**Supplementary figure 1. Schematic design of CFF measuring tool.** Custom CFF measuring tool from diagonal- (top left), top- (top right), front- (bottom left) and side view (bottom right). Red crossed circle indicates location of the LED light inside the housing. Blue dashed line indicates length of the viewing tube.

**S2: Supplementary analysis of RDM data**

We conducted a supplementary analysis on the RDM data to assess if the slight difference in methodologies between the experimenters caused significantly different task results. T-test indicated that the two data sets were not significantly different from each other (Welch's T = 0.28, degrees of freedom = 72.94, P = 0.778). Separate regression modelling also showed no significantly different effects of the predictors between the two data sets (Supplementary table 1).

**Supplementary table 1. Results of RDM analysis with separated dataset.** As the methodology of data collection of the experimenters differed slightly, we conducted a supplementary analysis in which the data collected through the two methodologies were treated as two separate data sets. *Top*: linear regression output of the part of the data set in which the Quest step was employed to titrate coherence levels during the task. *Bottom*: linear regression output of the part of the data set in which coherence levels were adjusted manually depending on results of the "Verify" step.

| **Model for Quest data**<br>**$R^2$ = -0.02** | | |
|---|---|---|
| **Predictor** | **Estimate (β)** | **P-value** |
| CFF | -0.11 | 0.753 |
| Age | -1.19 | 0.288 |
| Gender (male) | 6.33 | 0.313 |

| **Model for Verify data**<br>**$R^2$ = -0.10** | | |
|---|---|---|
| **Predictor** | **Estimate (β)** | **P-value** |
| CFF | -0.22 | 0.396 |
| Age | -2.02 | 0.352 |
| Gender (male) | 3.27 | 0.539 |
| Gender (other) | 1.61 | 0.896 |

**S3: Results of Roy-Bargmann stepdown analysis**

**Supplementary table 2. Roy-Bargmann stepdown models.** Post-hoc univariate models, testing our first priority response variable (AB magnitude) and our second priority response variable (overall T2 accuracy). P-values shown in bold indicate significance after Bonferroni correction (P < 0.0167). Note that in the model testing the second priority response variable, the first priority response variable is included as a covariate, as per the Roy-Bargmann stepdown procedure.

| **AB magnitude<br>Model $r^2$ = 0.079** | | |
|---|---|---|
| **Predictor** | **Estimate (β)** | **P-value** |
| CFF | -0.01 | **0.014** |
| Age | 0.02 | 0.066 |
| Gender (male) | -0.02 | 0.636 |
| Gender (other) | -0.14 | 0.066 |

| **Overall T2 accuracy<br>Model $r^2$ = 0.338** | | |
|---|---|---|
| **Predictor** | **Estimate (β)** | **P-value** |
| CFF | 0.01 | 0.503 |
| Age | -0.05 | 0.354 |
| Gender (male) | 0.83 | **0.002** |
| Gender (other) | 0.24 | 0.738 |
| AB magnitude | -3.13 | **$7.48^{-6}$** |
